# NetCrafter: Ontology-derived Gene Network Modeling and Functional Interpretation

**DOI:** 10.64898/2026.01.16.699831

**Authors:** Yeji Lee, Soyeong Kim, Yuna Park, Eunah Jeong, Sumin Jeong, Seyeon Kim, Jaemoon Shin, Euna Jeong, Hohsuk Noh, Sukjoon Yoon

## Abstract

Understanding the complex nature of multi-functional interactions among genes is crucial for interpreting omics data. We developed NetCrafter, an ontology-driven platform for constructing *de novo* gene networks that are specific to each input gene list and quantitatively defined by ontology-weighted similarity. By incorporating the probabilistic association of ontology or curated gene sets into a weighted Tanimoto similarity metric, NetCrafter transforms enrichment results into quantitative semantic similarity scores between genes, enabling the creation of context-specific statistical networks. These networks can be further decomposed into optimal sub-networks, facilitating multi-functional interpretation and the identification of gene interaction hotspots. NetCrafter also supports the integration of heterogeneous omics–derived gene lists through consensus ontology scoring. Importantly, this list-specific, quantitative framework reveals functional hotspots and target-biomarker relationships – even in cases where ontology terms alone are not predictive of node-level attributes such as CRISPR efficacy. NetCrafter provides an interactive platform for constructing and interpreting dynamic, context-specific gene networks, leveraging ontology-based functional associations to uncover underlying mechanisms and identify key nodes. It is freely available at https://netcrafter.sookmyung.ac.kr and integrated into Q-omics platform (https://qomics.ai) to enhance the utility of cancer omics data.

## Introduction

Constructing gene networks has emerged as a powerful approach to visualize and analyze complex multi-functional interactions among genes, providing insights into underlying biological processes, functional relationships, and phenotypes[1–3]. However, most existing tools for gene network construction rely on predefined templates[4–6], fixed ontological frameworks[7–10], or specific data types[11, 12]. These limitations constrain their ability to generate networks that are specific to the input data and quantitatively defined, thereby limiting applications in complex omics analyses. In practice, template-based or ontology-fixed networks often treat interactions as static and binary, making it difficult to recalibrate edge definitions for a new gene list or to compare heterogeneous layers such as RNA expression and CRISPR knockout profiles. An alternative approach is to re-evaluate existing ontological resources and gene set databases in a gene list–specific and quantitatively scalable manner, enabling unified analysis across multiple omics datasets and diverse ontologies. To overcome these limitations, ontologies can be treated as dynamic statistical resources whose contributions are recalibrated for each input gene list. This concept highlights the need for methods that dynamically rebuild functional relationships rather than relying on fixed templates — an ability that conventional frameworks do not provide. Recent advances have introduced new network-construction tools such as knowledge-integrative builders (e.g., NeKo[13]) and modern multi-omics or single-cell network-inference frameworks (e.g., DeepMAPS[14], SCRIPro[15], and MINIE[16]). While powerful within their respective domains, these approaches still rely on predefined correlations, prior knowledge, or model-specific assumptions, and therefore do not support de novo, ontology-weighted, list-specific network reconstruction.

To address these challenges, we present NetCrafter, a versatile tool for the *de novo* generation of ontology- and curated gene set-based quantitative networks to overlay omics data and facilitate their multi-functional interpretation. Omics data, such as RNA expression and CRISPR profile, are frequently analyzed within networks to elucidate gene-gene interactions represented in the data[17–19]. Once a specific set of genes is provided, NetCrafter calculates probabilistic associations of ontology terms (or functional signatures) with the gene list and incorporates them as weights into a similarity metric to quantify overlapping terms between genes. This approach enables the creation of context-specific, quantitative network frameworks that are scalable for hierarchical decomposition and flexible for multi-functional interpretation.

In this study, we integrated diverse ontology and gene signature databases such as GOBP (Gene Ontology Biological Process)[20], HPO (Human Phenotype Ontology)[21], Hallmark gene sets [22, 23] and Oncogenic Signature collections [22, 23] to derive semantic similarity between genes. The *p-value* of each functional term for a given gene list was used as a weight to summarize all overlapping terms between genes. The weighted Tanimoto score was then used to define quantitative edges, enabling the construction of customizable gene networks that overcome the limitations of fixed function-gene mappings. These networks were flexibly decomposed and interpreted at varying thresholds of weighted Tanimoto scores. Previous studies have reported that hub nodes in biological networks play essential roles in interpreting the overlaid data[24, 25]. Here, we investigate how the distribution of weighted Tanimoto scores correlates with network hotspots, such as drug targets.

NetCrafter provides user-friendly, interactive tools for analyzing and interpreting consensus functions and phenotypes across varying levels of statistical confidence. Furthermore, its integration with the Q-omics platform[26, 27] enhances its utility for cancer research by enabling seamless interpretation of diverse omics datasets. This report highlights the innovations of NetCrafter and demonstrates its ability to uncover functional insights and patterns in omics data. Therefore, in this study, we demonstrate NetCrafter using two representative gene lists, illustrating how it constructs quantitative functional networks, decomposes subnetworks, and integrates multi-source datasets. The following sections illustrate how these capabilities reveal functional hotspots, consensus modules, and actionable target–biomarker relationships.

## Results

### Gene network construction and ontology clustering

Ontology-based semantic similarity between genes was quantified to create unique networks for a given gene list (Figure 1A). A total of 7,172 functional ontologies were retrieved from GOBP terms, each containing 3 to 300 genes (https://geneontology.org/). Similarly, 7,342 HPO terms (https://hpo.jax.org/), each containing 3 to 300 genes, were included to represent disease phenotypes. The association *p-value* of ontology terms with the given gene list was calculated using Fisher’s exact test[28, 29]. The shared ontology terms between gene G1 and G2 were quantified using the weighted Tanimoto index (*Tw(G1,G2)*), calculated based on the -*log(p-value)*s of all overlapping ontology terms. The inverse of *Tw(G1,G2)* was used to define the edge length between genes. For each gene pair in the list, edges satisfying a *Tw(G1,G2)* cutoff were used to construct discrete networks, which were visualized using a force-directed layout algorithm[30]. Node size reflects either the sum of *Tw* from directly connected edges, the sum of *-log(p-value)* of associated terms or the corresponding omics data.

**Figure 1.**
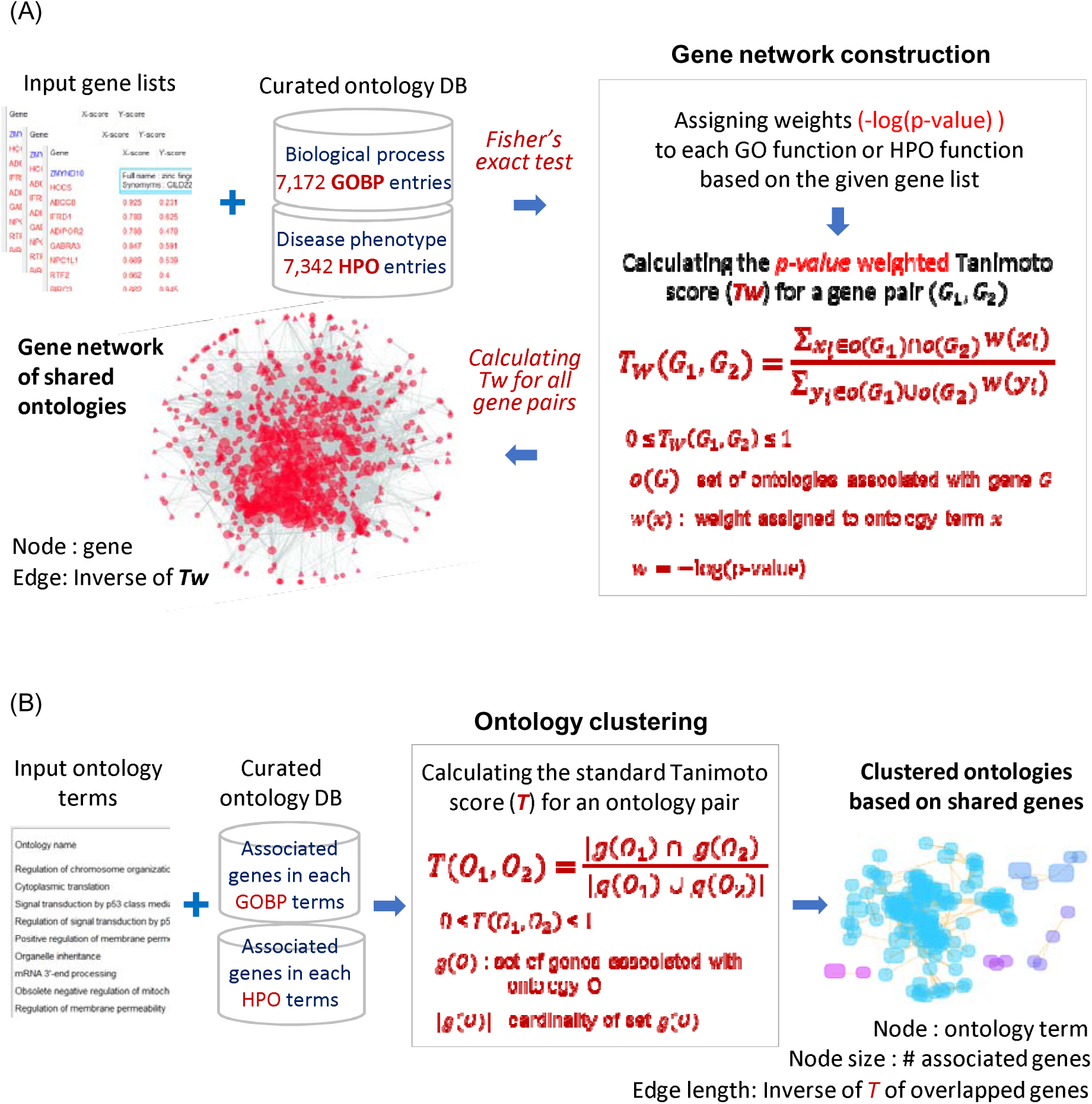
Workflow of gene network construction and ontology cluttering. The figure Illustrates the process of generating networks using Gene Ontology Biological Process (GOBP) terms and Human Phenotype Ontology (HPO) terms containing 3 to 300 genes. (A) Creating gene networks of shared ontologies: For a gene list, the -*log*(*p-value*) of each ontology term was calculated using Fisher’s exact test. *Tw*(*G*_1_,*G*_2_) denotes the *p-value* weighted Tanimoto score. The *p-value* combination method for multiple gene lists is described in the Methods section. (B) Ontology clustering based on shared genes: Input oncology terms were clustered using the regular Tanimoto Index between terms (*T*(*O*_1,_*O*_2_), which measures the overlap in shared genes.

Associated ontology terms for a gene list or gene network, were clustered into networks using the regular Tanimoto index (*T(O1,O2)*), which summarizes overlapping genes between ontology term O1 and O2 (Figure 1B). These networks display clustered patterns of ontology terms based on shared genes. Given the extensive number of functions included in a gene network, identifying the representative ontology terms contributing to the network can be challenging. In NetCrafter, representative ontology terms are quantitatively defined based on clustering criteria such as the -*log(p-value)* or the number of associated genes.

### Analysis of weighted Tanimoto (Tw) scores in gene networks

The semantic similarity between genes was quantified using the probabilistic sum of all overlapping ontology terms, denoted as *Tw* (Figure 1A). With the extensive coverage of 7,172 GO terms, 7,342 HPO terms and 189 oncogenic signatures (with more to come), most genes share one or more functional terms, resulting in their connection by edges with a wide range of *Tw* values. Networks with higher semantic similarity were defined by applying stricter *Tw* cutoffs. To determine a reasonable cutoff level for *Tw*, we simulated the distribution of the *Tw* scores in GOBP-based networks using randomly selected gene lists of varying sizes (Supplementary Figure 1A). As the gene list size increased, the top 1%, 5%, 10% and 15% cutoffs were slightly decreased (Supplementary Figure 1B). Average *Tw* cutoffs across gene lists of varying sizes, were subsequently used in network analysis to enhance statistical significance (Table 1). Comparatively, HPO terms with lower gene coverage than GOBP, exhibited lower *Tw* scores (Supplementary Figure 1C and 1D). Based on these *Tw* distributions, we defined top 1%, 5%, 10% and 15% cutoffs to provide statistical confidence for defining discrete subnetworks in NetCrafter. For ontology clustering (Supplement Figure 2), the distribution of *T* scores was also simulated, and statistical confidence levels were established for functional interpretation (Table 1).

**Table 1.** Average *Tw* and *T* cutoffs for gene networks and ontology clusters. Average Tw and average T indicate the mean (weighted) Tanimoto values of connected edges. The average *Tw* values are derived from gene lists ranging in size from 10 to 1,000 as shown in Supplementary Figure 1B and 1D. The average *T* values are derived from ontology lists ranging in size from 10 to 1,000 as shown in Supplementary Figure 2B and 2D.

| Statistical confidence | Average $T_w$ of gene networks | | Average $T$ of ontology clustering | |
| --- | --- | --- | --- | --- |
|  | GOBP | HPO | GOBP | HPO |
| 0.01 (top 1%) | 0.58 | 0.26 | 0.146 | 0.204 |
| 0.05 (top 5%) | 0.30 | 0.16 | 0.086 | 0.116 |
| 0.10 (top 10%) | 0.23 | 0.12 | 0.068 | 0.090 |
| 0.15 (top 15%) | 0.19 | 0.10 | 0.057 | 0.076 |

### Decomposition of gene networks and interpretation

A total of 383 genes, whose CRISPR efficacy was significantly associated with TP53 mutations (*p<0.01*), were used for gene network construction (Figure 2A). A set of 1,720 GOBP functions shared within these genes contributed to the *Tw* scores and overall network structure (Figure 1A; see Methods for details). From these 1,720 terms, NetCrafter further selected 48 representative terms significantly associated with the 383 genes (default criteria: -*log(p-value*) > 2, total *Tw* > 1, shared genes (*T*) <0.3)) (Figure 2A). Users may adjust the threshold parameters to redefine the representative ontology terms. Among the selected terms in this example, “rRNA metabolic process” encompassing 18 genes, contributed the largest cumulative edge weight (total *Tw* = 131; red nodes in Figure 2A), whereas “Chromosome organization” encompassing 21 genes, was the most significantly enriched in the network (-*log(p-value*) = 5.7; blue nodes in Figure 2A). These subnetworks highlight functional hotspots in TP53-mutant contexts, suggesting potential vulnerabilities in ribosome biogenesis and chromosome organization. These TP53-associated subnetworks are biologically plausible, as perturbations in rRNA metabolic processes are known to trigger nucleolar stress, stabilizing TP53 via the ribosomal protein-MDM2 axis [31]. In addition, shared nodes such as *CENPE* and *LSM10* correspond to genes whose disruption induces chromosome instability or replication stress, consistent with their connectivity within the TP53-related functional modules. Specifically, *CENPE* is critical for chromosome organization, while *LSM10* is essential for histone mRNA processing. Defects in histone supply are known to trigger p53-dependent cell cycle arrest due to replication stress [32].

**Figure 2.**
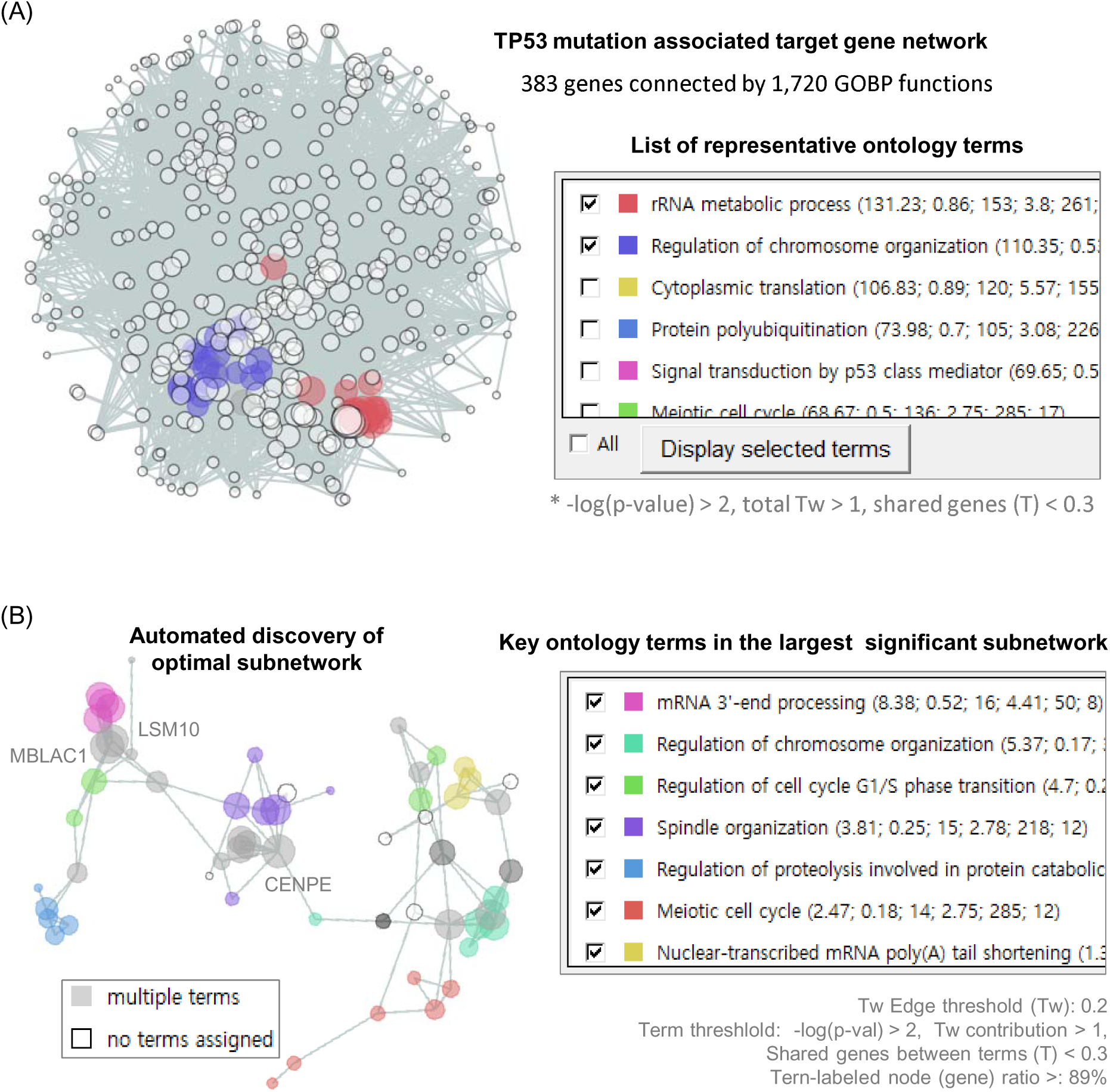
Decomposition of gene networks and functional analysis. A total of 383 genes whose CRISPR knockout efficacy was positively associated (*p*<0.01) with TP53 mutations in cancer cells were used for GOBP network construction and interpretation. (A) A total of 1,720 GOBP terms contributed to network edges with *Tw > 0*. Forty-eight representative GOBP terms were selected based on three criteria: significant term enrichment (p<0.01), sufficient edge contribution (total Tw > 1) and limited gene overlap between terms (T<0.3). (B) The automatically selected “Optimal subnet” highlights the distribution of 7 representative GO terms across 58 gene nodes. Three genes (MBLAC1, LSM10 and CENPE) were shared nodes between these representative terms.

Subsequently, users can decompose the network by increasing the *Tw* threshold to define more significantly connected subnetworks. NetCrafter automatically identifies and visualizes an optimal subnetwork by scanning the network structure across diverse *Tw* thresholds in combination with other criteria such as −log(*p-value*) of terms and the ratio of term-labeled nodes (Figure 2B). In the present demonstration, NetCrafter identified a subnetwork of 58 genes as the largest significant TP53 mutant-associated CRISPR target network. Seven terms were identified as the major contributing functions in this gene network. Three genes (MBLAC1, LSM10 and CENPE) served as shared nodes linking these key functions. This result illustrates how NetCrafter can pinpoint specific functional subnetworks and prioritize genes that bridge multiple processes, providing mechanistic insights into TP53-driven vulnerabilities. Users may further define subnetworks of varying size and connectivity by adjusting the threshold parameters.

Overall, this analysis demonstrates that at low *T*w thresholds, the gene network preserves a broad range of functional interactions, capturing genes and relationships that extend beyond those explicitly annotated under specific functional terms. By progressively increasing the threshold, subnetworks highlight shared genes within defined representative functional terms, enabling a more detailed understanding of gene interactions in multifunctional contexts. A similar NetCrafter analysis of TP53 target networks on shared “Oncogenic Signatures” is demonstrated in Supplementary Figure 3.

### Analysis of hierarchical network structure

The distributions of *Tw* scores were used to decompose networks and define discrete subnetworks with higher statistical confidence. As shown in Table 1, most genes are covered and remain analyzable in networks with *Tw* < 0.5 (Figure 3A). As the *Tw* threshold increases, the −log(*p-*value) distribution for associated GOBP terms changes slightly, although the number of associated terms decreased dramatically (Figure 3B). This suggests that many GOBP terms with low statistical significance (i.e., lower −log(*p-value*)) still contribute meaningfully to *Tw* scores. For instance, a high *Tw* score may result when two genes share numerous ontology terms, even if those terms have relatively low enrichment significance. In the TP53 subnetworks constructed using *Tw* > 0.3 (top 5% edges; Supplementary Table 1), the representative functions of subnetworks showed varied −log(*p-*value) scores (Figure 3C). In many cases, the dominant term – defined by the largest gene coverage – had a lower - log(*p*-value) and differed from the representative term with strongest enrichment (see Supplementary Table 1). For example, 4 genes involved in a signaling pathway were connected via *Tw* > 0.3 edges, despite being associated with 10 GOBP terms of relatively low significance (purple circle Figure 3C; also see Supplementary Table 1). Figure 3C also shows that the network size does not correlate with the number of associated GOBP terms. In the analysis of red subnetwork which includes 14 terms (Figure 3C and Supplementary Table 1), two terms – the dominant term, “Cytoplasmic translation” and the representative term, “rRNA metabolic process” - were associated with 7 and 6 genes, respectively (Figure 3D). The gene RPL7 served as a connecting point between the gene sets covered by these two terms.

**Figure 3.**
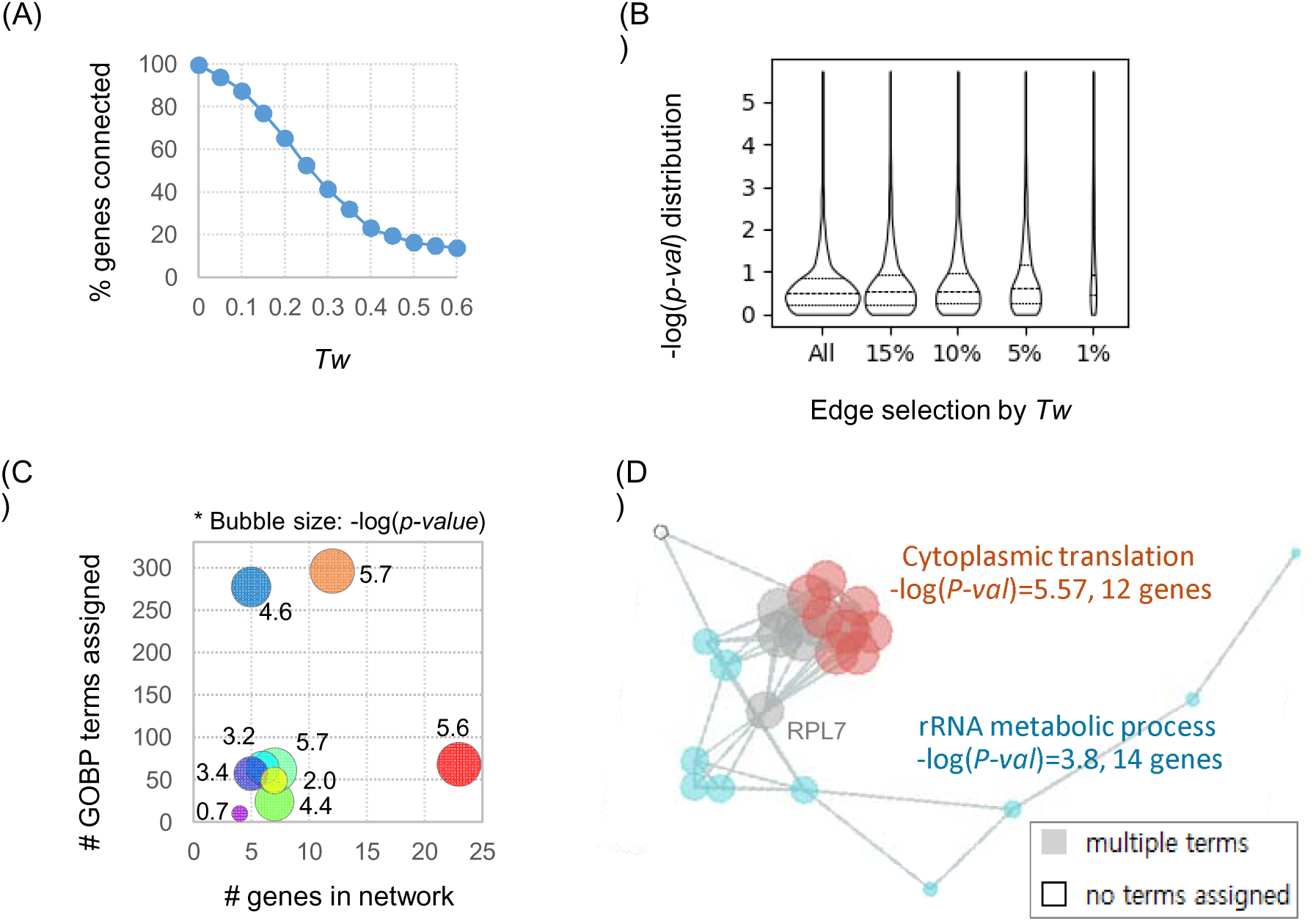
Contribution of enrichment score (*p-value*) versus consensus (i.e., *Tw*) to the hierarchical decomposition of TP53 target networks. (A) Changes in gene coverage during hierarchical decomposition of GOBP networks using *Tw* thresholds. (B) Distribution of the Fisher enrichment scores (-log(*p*-value)) of GOBP terms associated with the gene list during network decomposition by *Tw* threshold. (C) Comparison of network size and number of associated functions in TP53 target networks of *Tw* > 0.3 (Supplementary Table 1). The - log(*p*-value) of the representative function is shown numerically and also represented by bubble size. (D) Functional interpretation of a subnetwork in panel (C) (see also Supplementary Table 1).

### Network structure versus node attributes

We investigated whether ontology-based quantitative network structures have potential for prioritizing essential nodes based on their attributes. CRISPR knockout data for 383 genes (used in Figure 2) were analyzed as node attributes. The “sum of *-log(p-value)*” of associated GOBP terms for each gene node showed no correlation with CRISPR efficacy data (Figure 4A).

**Figure 4.**
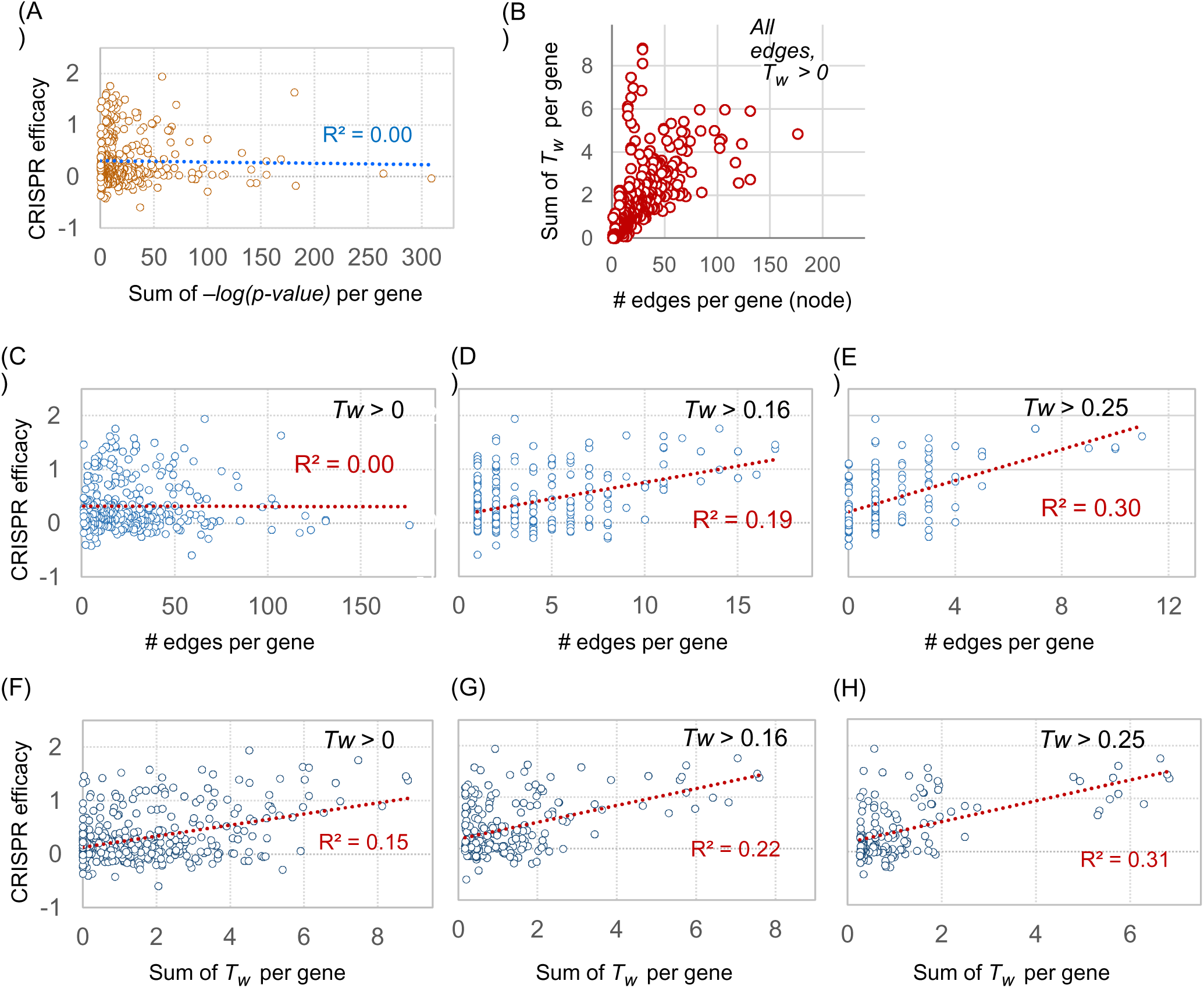
Comparison of ontology-based network structure with node attributes. CRISPR knockout efficacy data for 383 genes whose responses were associated with TP53 mutation status across diverse cancer lineages (same gene set as in Figure 2) were analyzed. (A) Comparison of the sum of *-log(p-value)* with CRISPR efficacy data for each gene node. (B) Relationship between the sum of *Tw* scores and the number of edges connected to each gene node in the TP53 gene network. (C-E) Comparison of the number of edges per node with CRISPR efficacy across different network configurations. (F-H) Comparison of the sum of *Tw* per node with CRISPR efficacy.

To explore the contribution of *Tw* to network structure, we analyzed the correlation between the number of edges and the sum of *Tw* scores for each node (Figure 4B). The sum of *Tw* scores was not directly correlated with the number of edges. Notably, some nodes with relatively few edges still exhibited high “sum of *Tw*” scores, suggesting that both the number of edges and the statistical confidence measure (sum of *Tw*) are essential for interpreting network structure.

The number of edges per node showed a correlation with CRISPR data only when edges were defined using higher *Tw* thresholds (Figure 4C-E). Edges with low statistical confidence did not support the prioritization of key nodes as CRISPR targets. Interestingly, when edge quantification was based on the sum of *Tw* scores per gene, correlations with CRISPR efficacy were observed even at low threshold (*Tw* > 0) (Figure 4F-H).

This analysis highlights that the network structure, as defined by edge distribution, has significant potential for prioritizing key nodes. Moreover, the quantitative measurement of edges using *Tw* scores offers enhanced resolution for identifying CRISPR targets. Beyond the analysis of 383 TP53 mutant associated genes, we also constructed diverse gene networks from CRISPR-based gene lists associated with various drugs and tumor suppressor genes across 20 tumor lineages (Figure 5). For each of these networks, the sum of Tw per gene was compared with experimentally measured CRISPR efficacy or drug response, and genes showing positive or negative associations with these functional readouts (red and blue points, respectively) consistently tended to occupy regions with higher Tw values. These results indicate that Tw-based network connectivity captures functionally important nodes across multiple independent perturbation datasets, supporting the general applicability of Tw-based node prioritization beyond the TP53 example.

**Figure 5.**
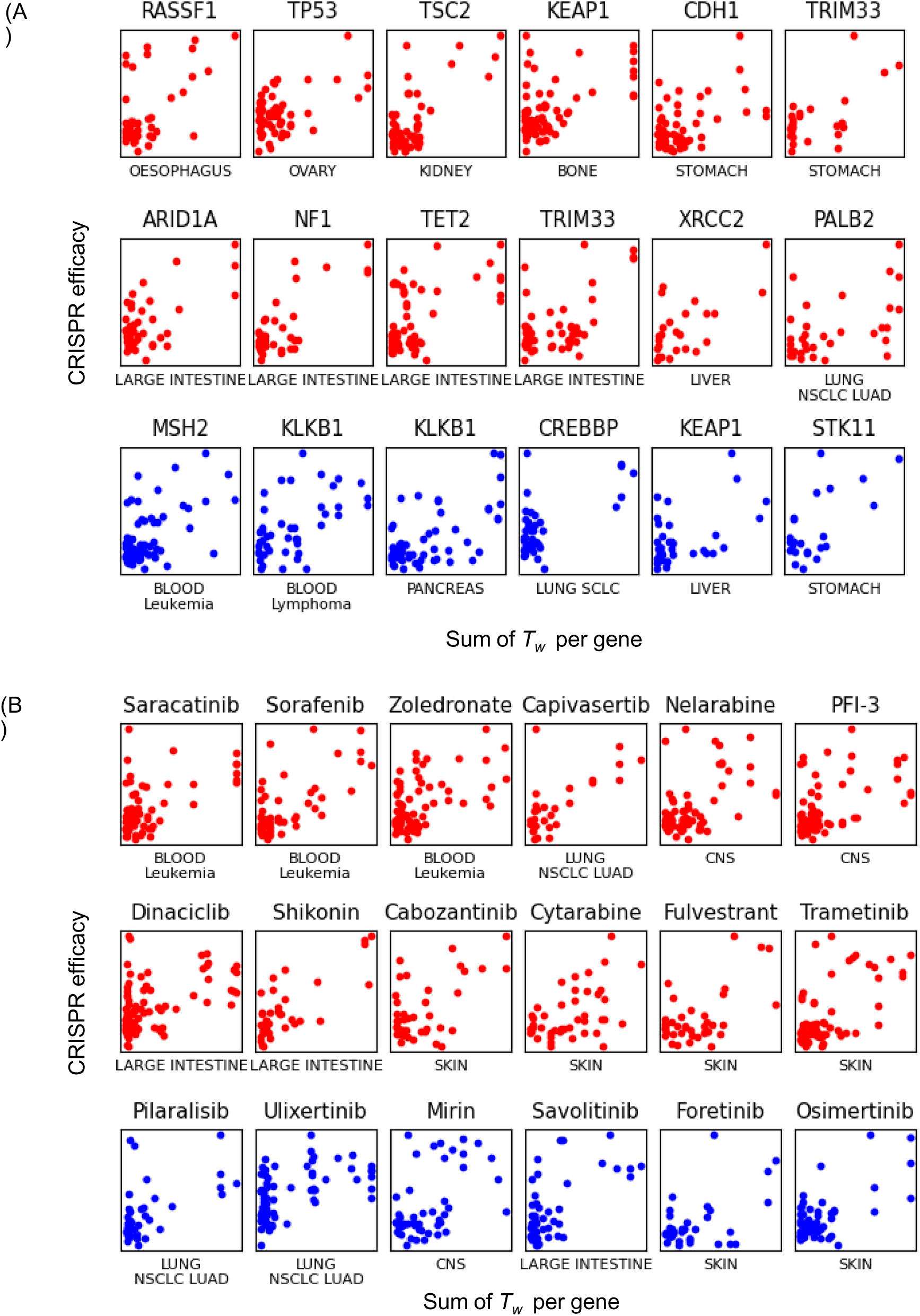
Comparison of ontology-based network structure with node attributes. CRISPR data for genes whose responses are significantly associated with (A) RNA expression of a tumor suppressor gene and (B) anticancer drug response across diverse cancer lineages, as retrieved from Q-omics. For each node in the gene networks, the sum of Tw was compared with CRISPR efficacy measurements. Red points represent genes whose CRISPR data show a positive correlation with tumor suppressor expression or drug response, and blue points represent genes with negative correlations.

### Meta-analysis of consensus gene networks derived from multiple omics datasets

NetCrafter provides a versatile platform for integrating independent omics datasets into ontology-based gene networks, enabling comprehensive meta-analysis and detailed subnetwork exploration. To support this, we calculated consensus −log(*p*-value) weight for ontology terms across multiple gene lists using *p*-value combination methods (See details in Methods section).

In this demonstration, 702 genes with RNA expression or CRISPR knockout data associated with TP53 mutant samples were used to create the network (Figure 6). The term “Mitotic nuclear division” (-log(*p*-value)=4.18) encompassed 12 genes (black circles), whereas the representative term, “Cytoplasmic translation” (-log(*p*-value)=5.57) covered only 11 genes. A total of seven subnetworks with high statistical confidence were generated using a threshold of *Tw* > 0.58 (top 1%) and node count > 3. Genes associated with “Mitotic nuclear division” were separated into two subnetworks, containing both RNA and CRISPR hits, thus illustrating potential target-biomarker relationships in TP53 mutant cells. Notably, NetCrafter revealed candidate CRISPR targets such as CDK5RAP2, MZT1, and BCCIP that were tightly connected with mitotic regulators (TTK, TRIP13, BUB1B, SPC25, RACGAP1, PRC1, TPX2, MYBL2, and MISP), highlighting them as previously underappreciated targets and biomarkers in TP53-mutant cancers. Two additional subnetworks associated with “Cytoplasmic translation” consisted exclusively only RNA expression hits, while three other subnetworks – linked to “DNA unwinding”, “Keratinization” and “Immune response” – primarily included CRISPR hits.

**Figure 6.**
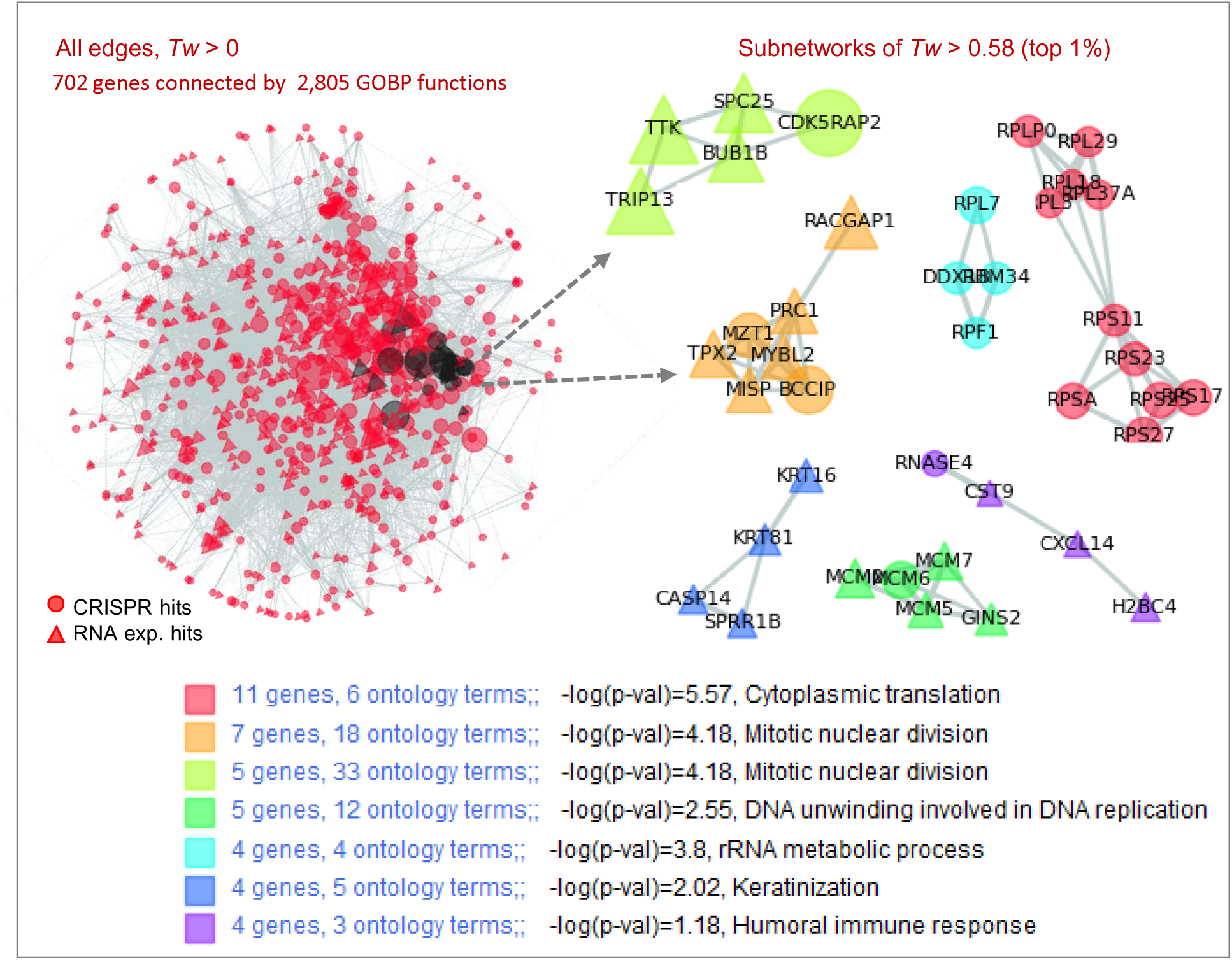
Consensus networks of TP53 mutant-associated CRISPR efficacy and RNA expression data. Gene networks were constructed using 385 CRISPR and 328 RNA datasets significantly associated (*p*<0.01) with TP53 mutations in caner cells. A total of 2,805 GOBP terms were linked to these genes. Seven subnetworks with *Tw* > 0.58 and node count > 3, are displayed with detailed descriptions provided below. Black-colored nodes indicated the 35 genes associated with the representative function “Mitotic nuclear division”. The subnetworks that include these genes are indicated with arrows.

This analysis provides a comprehensive view of the distribution of RNA and CRISPR hits across functional subnetworks, offering insight into the interaction between therapeutic targets (CRISPR) and biomarkers (RNA expression), supported by strong statistical confidence (top 1% edges).

## Discussion

Ontology- and curated signature-based gene networks provide a promising framework for the functional interpretation of omics data. Existing network tools often rely on pre-defined, static relationships or provide only categorical enrichment results, which limit their capacity to capture the quantitative and context-specific nature of gene interactions. In contrast, NetCrafter introduces a list-specific and quantitative framework in which gene sets are re-evaluated through Fisher’s exact test and transformed into weighted Tanimoto similarity scores. This converts enrichment outcomes into gene–gene similarity values that are both context-dependent and numerically defined, enabling dynamic, input-specific networks. Consequently, NetCrafter can be applied to gene lists derived from diverse biological contexts, including perturbation-derived CRISPR and drug-response datasets across multiple tumor lineages (Figure 5). Because it reconstructs networks dynamically for any input gene list, its applicability is not restricted to the cancer types used in our case studies and can be further extended to additional tumor types, multi-omics datasets, and future single-cell–derived gene sets. **Recent studies have also emphasized the importance of integrative data analysis and genomic resources in diverse tumor-specific contexts [33, 34], supporting the utility of NetCrafter for broader oncological applications.**

Genes interact within intricate contexts of overlapping functions, presenting challenges in omics interpretation. NetCrafter addresses this by automatically constructing decomposable networks based on statistical hierarchies of quantified functional similarity. Using two simple parameters - the statistical confidence of ontology-to-gene associations (-*log(p-value)*) and the probabilistic metric of gene-gene associations (*Tw*) (Figure 1) – it generates scalable networks that can be flexibly decomposed and interpreted. Additionally, the regular Tanimoto score (*T*) quantifies ontology-to-ontology similarity, allowing integrated analysis across diverse ontologies and gene signatures such as GOBP and HPO (Figure 2, 6 and Supplementary Figure 3).

A novel property of weighted Tanimoto gene network was observed, where network structures quantified by *Tw* scores demonstrated the potential to prioritize key nodes (Figure 4 and 5). While the significance of associated ontology terms themselves was not directly correlated with omics data attributes, such as CRISPR efficacy (Figure 4A), the sum of *Tw* values for a node showed emergent correlations with node attributes (Figure 4F-H). By leveraging extensive ontology information, NetCrafter generates a scalable *n2-*dimensional dataset of gene-gene relationships (edges), complementing conventional omics data and enabling graph-based modeling. Demonstrations in Figure 4 and Supplementary Figure 4 highlight its capability to identify functional hotspots and target-biomarker relationships.

In conclusion, NetCrafter establishes a new paradigm for ontology- and gene set-based network analysis by providing list-specific, quantitative and decomposable functional gene networks. It extends the utility of the Q-omics data mining platform through interactive visualization and automated interpretation, offering a broadly applicable tool for biological research and drug discovery. By uncovering TP53-associated functional hotspots (Figures 2–3) and novel candidate targets such as CDK5RAP2, MZT1, and BCCIP within target–biomarker subnetworks (Figure 6), NetCrafter demonstrates not only methodological advances but also the capacity to deliver new biological insights. These findings underscore its value as a discovery platform with translational potential for cancer biology and precision medicine.

A concise functional comparison with GeneMANIA[6], WGCNA[12], and widely used Cytoscape[35] ontology-based tools is summarized in Supplementary Table 2, highlighting that NetCrafter focuses on constructing ontology-weighted, list-specific, and decomposable gene networks rather than template- or overlap-based structures.

## Methods

### Availability and platform support

The NetCrafter software is available at https://netcrafter.sookmyung.ac.kr and is integrated into the Q-omics platform (https://qomics.io) to enhance the utility of cancer omics data. The software supports both Windows and macOS operating systems. Demo data used in this study are included in the “Demo_files” folder provided with the software installation. A comprehensive tutorial is also available on the NetCrafter homepage to assist users in utilizing its features.

### Data files for network construction

NetCrafter supports input files in CSV (Comma-Separated Values) format. Several examples of input files are provided in the “Demo_files” folder. The structure of these files is as follows:

- First row: Headers indicating the type of data
- Data rows (starting from the second row): Three columns in the order of:

1) Gene symbols: Up to 1,000 entries are supported for network construction. If a user provides more than 1,000 genes, NetCrafter analyzes up to 1,000 user-selected genes to maintain stable run time and memory usage.
2) Node types: Optional, ranging from 1 to 5. These values determine the shape of the nodes in the network visualization (See demo in Figure 6). If no value is provided, the default node type is set to 1, which corresponds to a circle shape.
3) Node data: Optional, such as RNA expression or CRISPR knockout data. This value determines the size of nodes in the network. If node data are not provided, the default node size is based on the number of associated ontologies.

### Gene networks and ontology clustering

Prior to network construction, weighted Tanimoto similarity (Tw) values were computed for all gene pairs using the equation shown in Figure 1A. Ontology-term enrichment p-values were obtained using Fisher’s exact test, and each term was assigned a weight of –log(p-value). In this framework, the Fisher p-values are used as weighting factors in the Tw similarity metric rather than as standalone hypothesis tests, so formal multiple-testing corrections (e.g., FDR or Bonferroni) are not applied to individual ontology terms. For each gene pair, the associated ontology terms and their intersection were identified, and the weighted overlap and union were calculated accordingly to derive Tw. These Tw values were then used as edge weights, and user-defined Tw thresholds were applied to construct subnetworks. For ontology-to-ontology similarity, the normal Tanimoto score shown in Figure 1B was used to cluster ontology terms and to remove redundant terms. The “average Tw” and “average T” refer to the mean (weighted) Tanimoto values of edges connected to gene nodes and ontology nodes, respectively.

To identify optimal subnetworks, Tw thresholds were increased in increments of 0.05. At each threshold, representative ontology terms were selected based on enrichment (–log(p-value) > 2), Tw contribution (> 1), and low redundancy (term-to-term Tanimoto < 0.3). Tw contribution was defined as the relative contribution of each ontology term to the weighted Tanimoto score, calculated as the weight of the term divided by the sum of the weights of all shared terms. Subnetworks with a term-labeled node ratio (i.e., the proportion of genes retained after term filtering) greater than 80% and more than one representative term were considered optimal.

NetCrafter generates gene networks based on either GOBP or HPO ontologies. By default, all edges with *Tw* > 0 are included. Users can interactively adjust the *Tw* threshold to display subnetworks. Representative ontology terms for each network are listed below the network visualization window. Additional details about associated terms are accessible via sub-windows. Ontology clustering is also available for terms associated with the gene network through the sub-window. For workflows involving independent GOBP or HPO entries in CSV files prepared by users, ontology clustering is supported. In this case, the second and third columns of the input file are not used. The current version of NetCrafter does not support user-uploaded ontology or gene-set files and operates with the built-in ontology and signature collections.

### Meta-analysis of consensus gene networks derived from multiple gene lists

Up to five gene datasets can be used to construct combined networks, with distinct node symbols representing separate datasets. The node index (1 to 5) indicates the datasets to which a gene belongs in the input file. The file “Genes_TP53mut_vs_multi_omics.csv”, located in the “Demo_files” directory of the NetCrafter software, provides an example of how to upload multiple gene lists, using values 1 to 5 in the “node_index” column.

For each gene list, Fisher’s p-values of ontology terms are calculated separately. Network edges between genes from different lists are computed using their respective p-value lists. To generate consensus networks for multiple gene lists, consensus p-values for ontology terms are calculated using p-value combination methods[36, 37]. The combined p-value can be determined as follows:

Let *p*_1_,…, *p_n_* be the association p-values of ontology term x for n gene lists. The combined p-value, *p*^(*comb*)^ is given by:

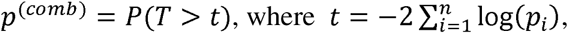

and *T* follows a chi-square (*χ*^2^) distribution with 2*n* degrees of freedom.

For the specific case of combining two gene lists, the following equality holds:

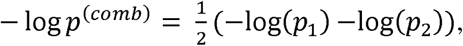

which simplifies the computation of Tw from the combined gene lists.

This consensus p-value procedure, based on Fisher’s combined probability test, helps reduce noise arising from multiple ontology-term evaluations.

### Preparation of TP53 mutant associated data for network analysis

A total of 385 genes whose CRISPR knockout results were significantly associated with TP53 mutations (*p-value < 0.01*) were retrieved from cell line data using Q-omics data mining. Additionally, 328 genes exhibiting significant RNA expression difference (*p-value < 0.01*) between TP53 mutant and wild type cancer patient samples were obtained through Q-omics. All other datasets used in the Supplementary figures were similarly acquired via Q-omics data mining.

## Acknowledgements

This work was supported by the National Research Foundation of Korea (NRF) Grant funded by the Korean Government (MSIP) [RS-2024-00345464 to Euna Jeong and RS-2023-00207857 to Sukjoon Yoon] and the Technology development Program funded by the Ministry of SMEs and Startups (MSS, Korea) [RS-2025-25403981 to Sukjoon Yoon].

## Author contributions

Soyeong Kim and Yuna Park - Analysis, Methodology, Software, Writing; Sumin Jeong - Analysis, Methodology; Seyeon Kim - Analysis, Software; Jaemoon Shin – Analysis, Methodology; Euna Jeong - Analysis, Methodology, Writing; Hohsuk Noh – Algorithm, Methodology; Sukjoon Yoon - Analysis, Conceptualization, Methodology, Supervision, Writing; All authors read and approved the final version.

## Key points

1) Creates unique gene networks using ontology-based weighted Tanimoto similarity scores
2) Enables hierarchical decomposition and flexible functional interpretation of gene networks
3) Supports the integration of multiple omics-derived gene lists through consensus ontology scoring
4) Identifies functional hotspots within gene networks

## Data Availability Statement

All data analyzed in this study are available through the NetCrafter and Q-omics platforms. The NetCrafter web server is freely accessible at https://netcrafter.sookmyung.ac.kr. Additional processed data are available from the corresponding author upon reasonable request.

