## Supplementary material for "NetCrafter: Ontology-derived Gene Network Modeling and Functional Interpretation": Supplment figures

### Slide 1
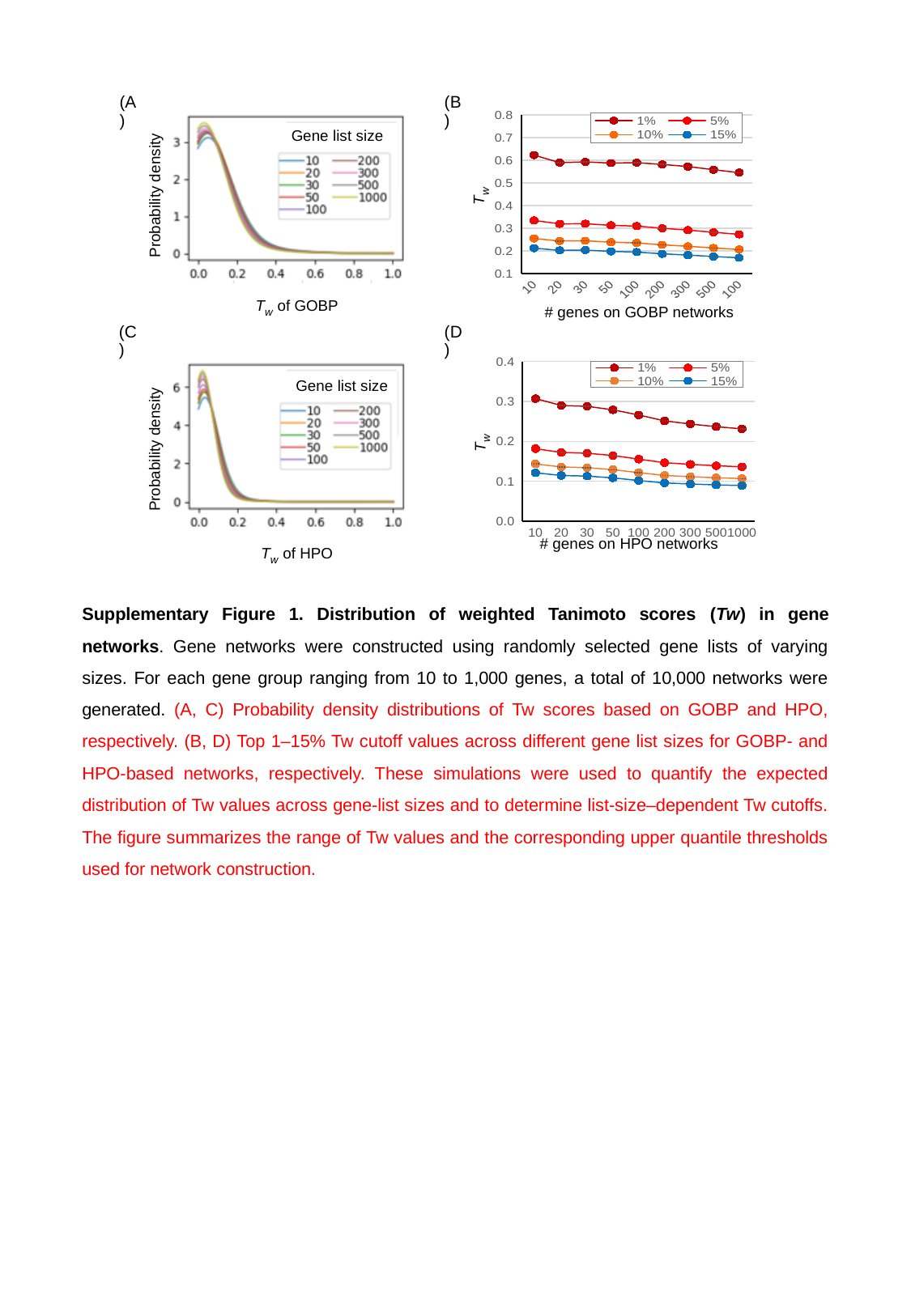

(A)
(B)
###### Chart
| Category | 1% | 5% | 10% | 15% |
|---|---|---|---|---|
| 10 | 0.623037647119235 | 0.334507212551385 | 0.254752654754496 | 0.211936667131448 |
| 20 | 0.589839971078292 | 0.319378221213566 | 0.243424333493533 | 0.202335472476641 |
| 30 | 0.593063341038507 | 0.320039087215639 | 0.244618773484981 | 0.203674508065856 |
| 50 | 0.587812160131375 | 0.312790501666171 | 0.238237129339115 | 0.197837479081522 |
| 100 | 0.589820304200095 | 0.309480331701745 | 0.235086859511568 | 0.194748830046262 |
| 200 | 0.581676103224 | 0.299799476133432 | 0.226692196475481 | 0.187042866802532 |
| 300 | 0.57249024701969 | 0.291830928845194 | 0.220246190981959 | 0.181415638503647 |
| 500 | 0.558652854131105 | 0.282063822437312 | 0.212794998320779 | 0.175124113755506 |
| 1000 | 0.545641997583609 | 0.273020962296438 | 0.206158572543059 | 0.16966657563372 |
 Gene list size
 Probability density
 T_w_
 T_w_ of GOBP
#### genes on GOBP networks
(C)
(D)
###### Chart
| Category | 1% | 5% | 10% | 15% |
|---|---|---|---|---|
| 10 | 0.306916579269606 | 0.181362711236983 | 0.143307862264055 | 0.121007121942028 |
| 20 | 0.28965279999972 | 0.171912606090155 | 0.135548130382653 | 0.114288101835059 |
| 30 | 0.287608724373109 | 0.170063456739113 | 0.133867063861795 | 0.112699326714657 |
| 50 | 0.278776846128955 | 0.164275607469185 | 0.1289519117432 | 0.108279804137279 |
| 100 | 0.2658035602488 | 0.155238077517857 | 0.121386581331189 | 0.101577951297362 |
| 200 | 0.251112952551149 | 0.146353047319369 | 0.114339646420328 | 0.0955621757432338 |
| 300 | 0.243298565573021 | 0.14224227030405 | 0.111204499188473 | 0.0929569659134738 |
| 500 | 0.236628539762509 | 0.138730702390717 | 0.108538077695653 | 0.0907535606959181 |
| 1000 | 0.231114183715407 | 0.13603305295361 | 0.106548796038601 | 0.089138574066053 |
 Gene list size
 T_w_
 Probability density
#### genes on HPO networks
 T_w_ of HPO
Supplementary Figure 1. Distribution of weighted Tanimoto scores (Tw) in gene networks. Gene networks were constructed using randomly selected gene lists of varying sizes. For each gene group ranging from 10 to 1,000 genes, a total of 10,000 networks were generated. (A, C) Probability density distributions of Tw scores based on GOBP and HPO, respectively. (B, D) Top 1–15% Tw cutoff values across different gene list sizes for GOBP- and HPO-based networks, respectively. These simulations were used to quantify the expected distribution of Tw values across gene-list sizes and to determine list-size–dependent Tw cutoffs. The figure summarizes the range of Tw values and the corresponding upper quantile thresholds used for network construction.

### Slide 2
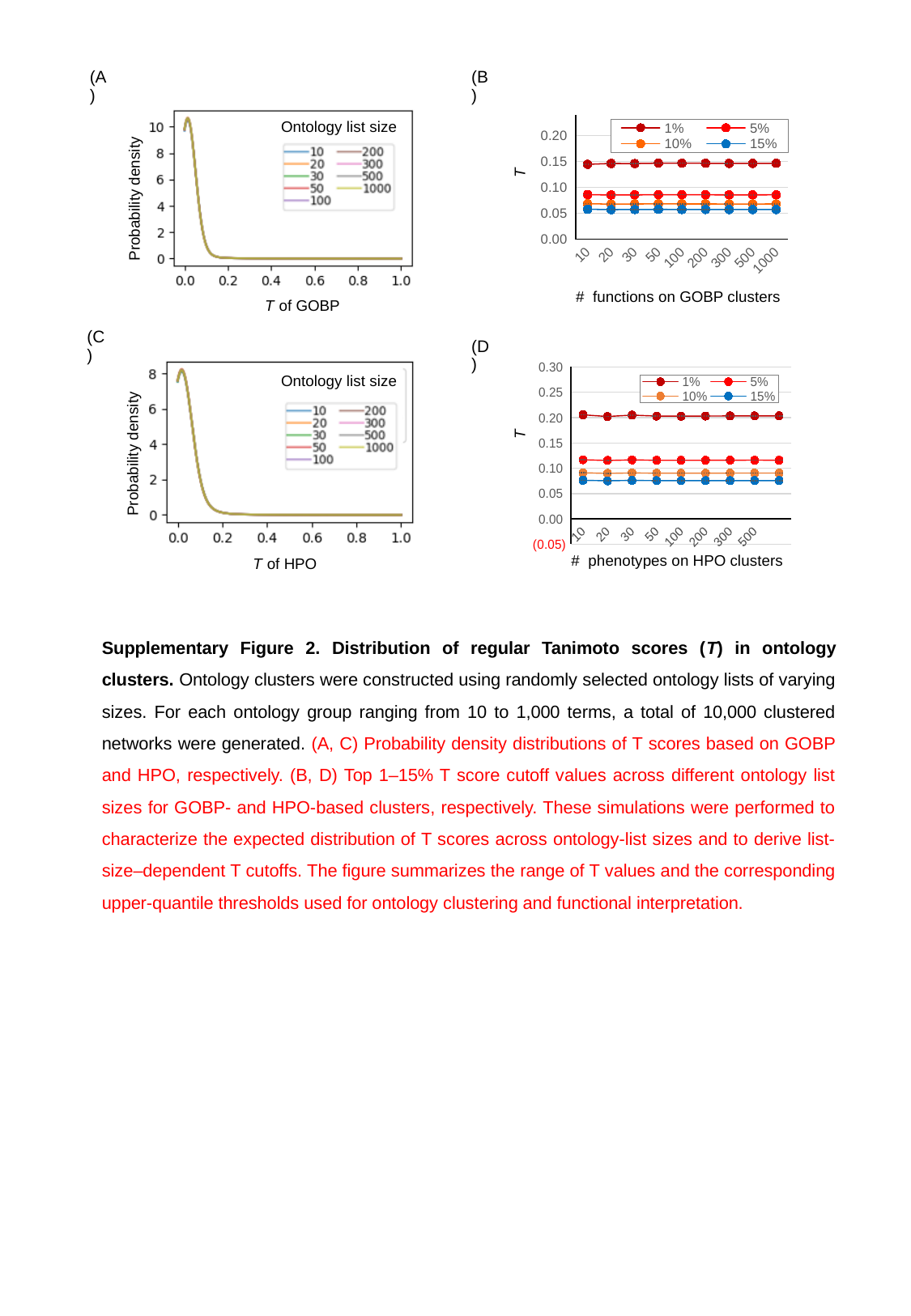

(A)
(B)
###### Chart
| Category | 1% | 5% | 10% | 15% |
|---|---|---|---|---|
| 10 | 0.144638501807665 | 0.0862805923043214 | 0.0685730241212523 | 0.0578331512615944 |
| 20 | 0.146305119462761 | 0.0853782410139395 | 0.0677416970588949 | 0.0571182028134839 |
| 30 | 0.14606937477881 | 0.0856501817473648 | 0.0679963595741051 | 0.0573406469554186 |
| 50 | 0.146849293591614 | 0.086029490389375 | 0.0682725774373512 | 0.0575652084389246 |
| 100 | 0.146628282807482 | 0.0858283115155454 | 0.0681044799855937 | 0.057420399806884 |
| 200 | 0.146653938551003 | 0.0858305448284447 | 0.0681111125788969 | 0.05742830498876 |
| 300 | 0.146365319837509 | 0.0856992128174819 | 0.0680127210790347 | 0.0573502890669938 |
| 500 | 0.146378912485202 | 0.0856706834420402 | 0.0679856108820057 | 0.0573255614773117 |
| 1000 | 0.146630572987485 | 0.0857576223863582 | 0.0680514374522641 | 0.0573794189575532 |
 Ontology list size
T
 Probability density
#### functions on GOBP clusters
 T of GOBP
(C)
(D)
###### Chart
| Category | 1% | 5% | 10% | 15% |
|---|---|---|---|---|
| 10 | 0.205551022511901 | 0.116713255166195 | 0.0910173892216132 | 0.076147446935302 |
| 20 | 0.202185410140195 | 0.115387258323646 | 0.089906394321857 | 0.075207245467238 |
| 30 | 0.20487353698857 | 0.11644133333598 | 0.0907946586362968 | 0.0759634067474217 |
| 50 | 0.202980433053644 | 0.115726265625489 | 0.0902186231174163 | 0.0754844884387773 |
| 100 | 0.202774552025505 | 0.115633155766982 | 0.0901503089759624 | 0.0754277080069541 |
| 200 | 0.203081324300259 | 0.115772348658943 | 0.0902432423465826 | 0.0754973755009476 |
| 300 | 0.203343689968388 | 0.115935404167373 | 0.0903846713531165 | 0.0756199452398471 |
| 500 | 0.203374228370615 | 0.115973432943689 | 0.090406533682452 | 0.0756350155873652 |
| 1000 | 0.20341720868948 | 0.115964671173341 | 0.0903960110523385 | 0.0756252211827308 |
 Ontology list size
T
 Probability density
#### phenotypes on HPO clusters
 T of HPO
Supplementary Figure 2. Distribution of regular Tanimoto scores (T) in ontology clusters. Ontology clusters were constructed using randomly selected ontology lists of varying sizes. For each ontology group ranging from 10 to 1,000 terms, a total of 10,000 clustered networks were generated. (A, C) Probability density distributions of T scores based on GOBP and HPO, respectively. (B, D) Top 1–15% T score cutoff values across different ontology list sizes for GOBP- and HPO-based clusters, respectively. These simulations were performed to characterize the expected distribution of T scores across ontology-list sizes and to derive list-size–dependent T cutoffs. The figure summarizes the range of T values and the corresponding upper-quantile thresholds used for ontology clustering and functional interpretation.

### Slide 3
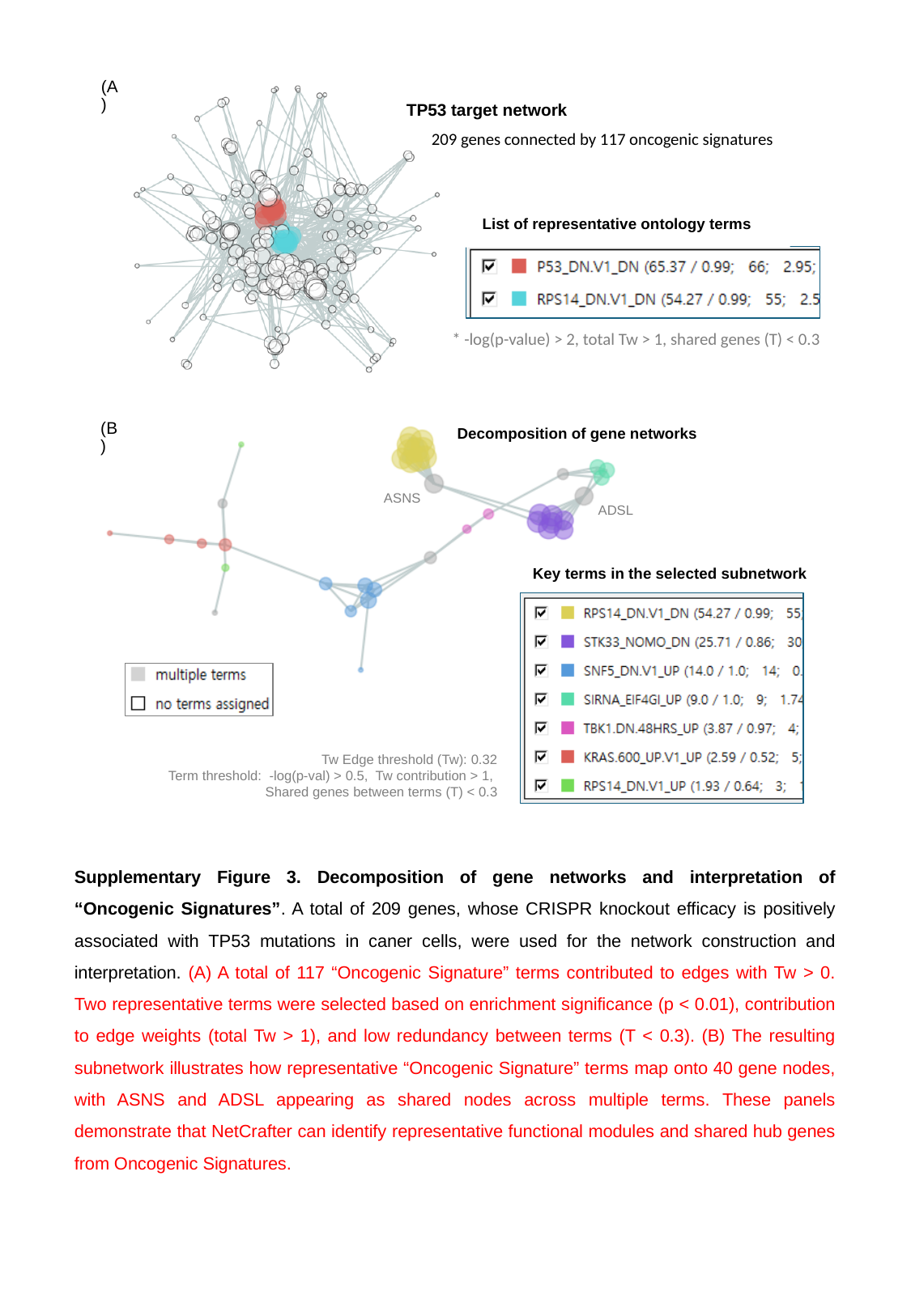

(A)
TP53 target network
209 genes connected by 117 oncogenic signatures
List of representative ontology terms
* -log(p-value) > 2, total Tw > 1, shared genes (T) < 0.3
Decomposition of gene networks
(B)
ASNS
ADSL
Key terms in the selected subnetwork
 Tw Edge threshold (Tw): 0.32
 Term threshold: -log(p-val) > 0.5, Tw contribution > 1,
 Shared genes between terms (T) < 0.3
Supplementary Figure 3. Decomposition of gene networks and interpretation of “Oncogenic Signatures”. A total of 209 genes, whose CRISPR knockout efficacy is positively associated with TP53 mutations in caner cells, were used for the network construction and interpretation. (A) A total of 117 “Oncogenic Signature” terms contributed to edges with Tw > 0. Two representative terms were selected based on enrichment significance (p < 0.01), contribution to edge weights (total Tw > 1), and low redundancy between terms (T < 0.3). (B) The resulting subnetwork illustrates how representative “Oncogenic Signature” terms map onto 40 gene nodes, with ASNS and ADSL appearing as shared nodes across multiple terms. These panels demonstrate that NetCrafter can identify representative functional modules and shared hub genes from Oncogenic Signatures.
