## Supplement table for "NetCrafter: Ontology-derived Gene Network Modeling and Functional Interpretation"

### Slide 1
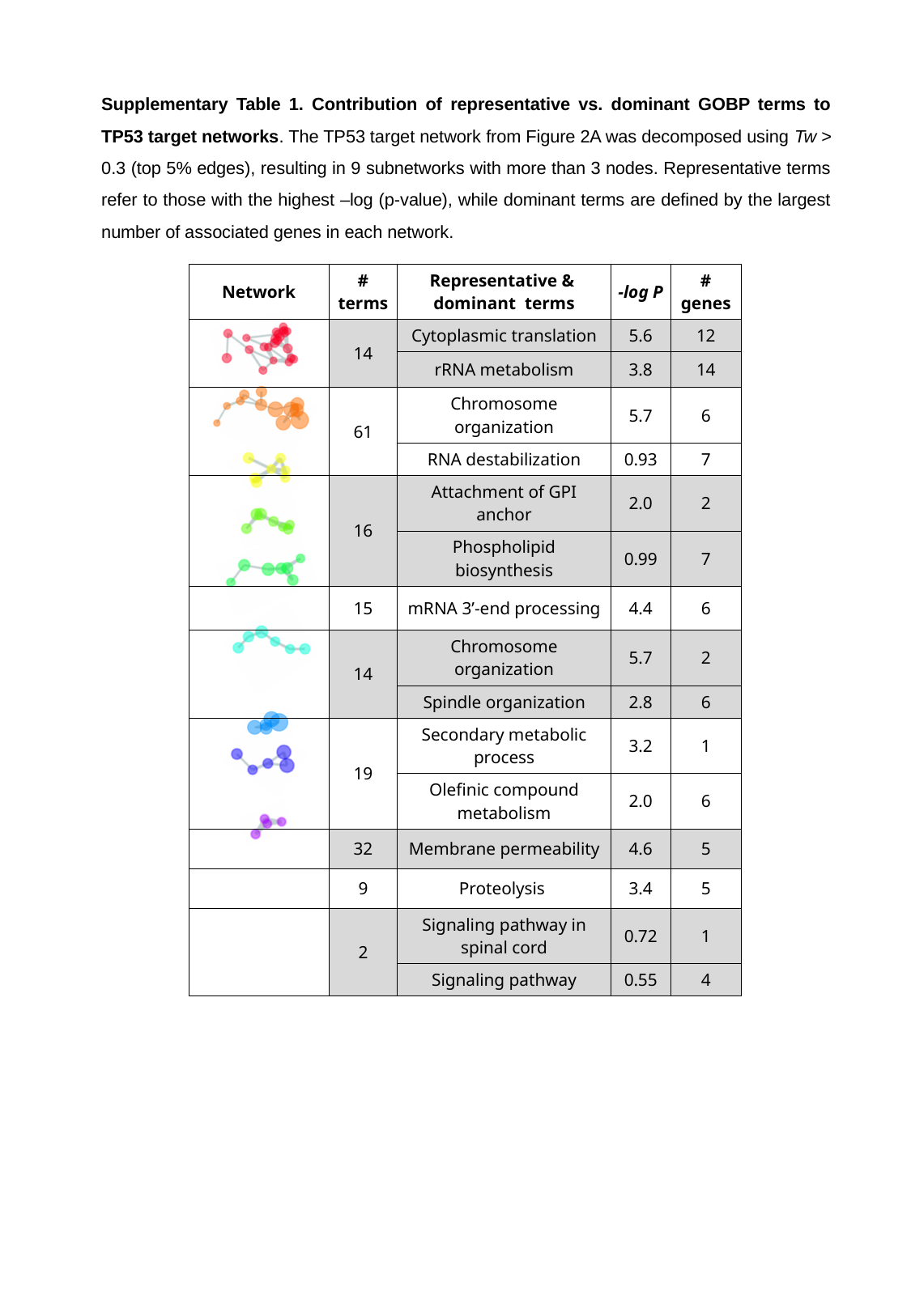

Supplementary Table 1. Contribution of representative vs. dominant GOBP terms to TP53 target networks. The TP53 target network from Figure 2A was decomposed using Tw > 0.3 (top 5% edges), resulting in 9 subnetworks with more than 3 nodes. Representative terms refer to those with the highest –log (p-value), while dominant terms are defined by the largest number of associated genes in each network.
| Network | # terms | Representative & dominant terms | -log P | # genes |
| --- | --- | --- | --- | --- |
| | 14 | Cytoplasmic translation | 5.6 | 12 |
| | | rRNA metabolism | 3.8 | 14 |
| | 61 | Chromosome organization | 5.7 | 6 |
| | | RNA destabilization | 0.93 | 7 |
| | 16 | Attachment of GPI anchor | 2.0 | 2 |
| | | Phospholipid biosynthesis | 0.99 | 7 |
| | 15 | mRNA 3’-end processing | 4.4 | 6 |
| | 14 | Chromosome organization | 5.7 | 2 |
| | | Spindle organization | 2.8 | 6 |
| | 19 | Secondary metabolic process | 3.2 | 1 |
| | | Olefinic compound metabolism | 2.0 | 6 |
| | 32 | Membrane permeability | 4.6 | 5 |
| | 9 | Proteolysis | 3.4 | 5 |
| | 2 | Signaling pathway in spinal cord | 0.72 | 1 |
| | | Signaling pathway | 0.55 | 4 |

### Slide 2
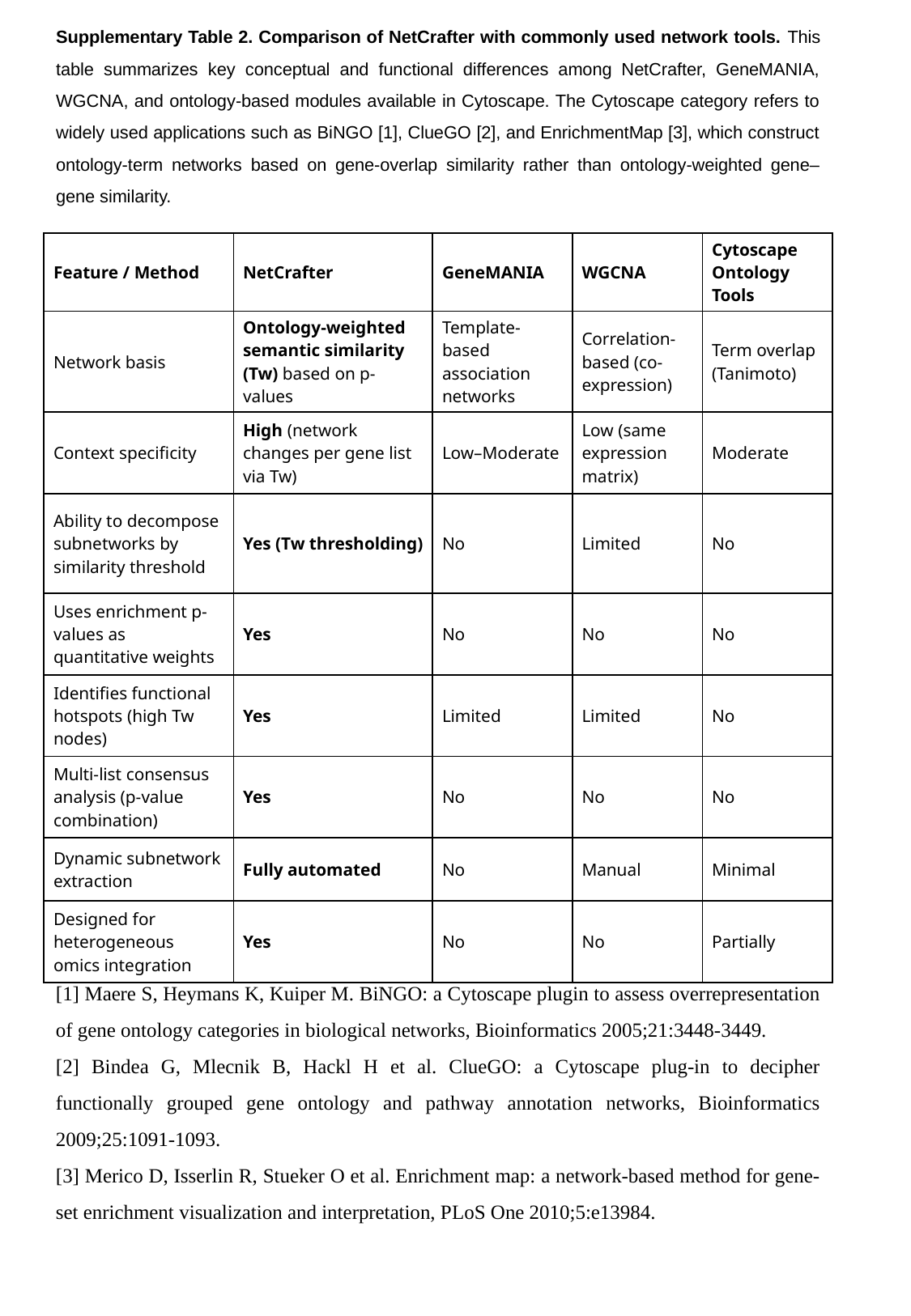

Supplementary Table 2. Comparison of NetCrafter with commonly used network tools. This table summarizes key conceptual and functional differences among NetCrafter, GeneMANIA, WGCNA, and ontology-based modules available in Cytoscape. The Cytoscape category refers to widely used applications such as BiNGO [1], ClueGO [2], and EnrichmentMap [3], which construct ontology-term networks based on gene-overlap similarity rather than ontology-weighted gene–gene similarity.
| Feature / Method | NetCrafter | GeneMANIA | WGCNA | Cytoscape Ontology Tools |
| --- | --- | --- | --- | --- |
| Network basis | Ontology-weighted semantic similarity (Tw) based on p-values | Template-based association networks | Correlation-based (co-expression) | Term overlap (Tanimoto) |
| Context specificity | High (network changes per gene list via Tw) | Low–Moderate | Low (same expression matrix) | Moderate |
| Ability to decompose subnetworks by similarity threshold | Yes (Tw thresholding) | No | Limited | No |
| Uses enrichment p-values as quantitative weights | Yes | No | No | No |
| Identifies functional hotspots (high Tw nodes) | Yes | Limited | Limited | No |
| Multi-list consensus analysis (p-value combination) | Yes | No | No | No |
| Dynamic subnetwork extraction | Fully automated | No | Manual | Minimal |
| Designed for heterogeneous omics integration | Yes | No | No | Partially |
[1] Maere S, Heymans K, Kuiper M. BiNGO: a Cytoscape plugin to assess overrepresentation of gene ontology categories in biological networks, Bioinformatics 2005;21:3448-3449.
[2] Bindea G, Mlecnik B, Hackl H et al. ClueGO: a Cytoscape plug-in to decipher functionally grouped gene ontology and pathway annotation networks, Bioinformatics 2009;25:1091-1093.
[3] Merico D, Isserlin R, Stueker O et al. Enrichment map: a network-based method for gene-set enrichment visualization and interpretation, PLoS One 2010;5:e13984.
